# CytoRipple: Mapping Tissue Polarization through Subcellular Cytoskeleton Dynamics

**DOI:** 10.64898/2026.10.13.755787

**Authors:** Daniel G. Chen, Taejus Yee, Justin Saco, J.J. Patten, Andreas I. Lackner, Michael Angelo, James R. Heath, Antoni Ribas, Katie M. Campbell

## Abstract

Spatial multi-omic technologies entail simultaneous measurement of tens-to-thousands of molecular features (e.g. RNA, protein). While cell type and state profiling is the major focus of spatial multi-omic analyses, cytoskeletal proteins are often solely utilized for cell segmentation and quality control. Here, we introduce a new computational framework, CytoRipple, that leverages these concomitant spatially-resolved single-cell measurements of cytoskeleton proteins to identify tissue-level vector fields. Polarization vectors are known to associate with cell movement and phenotype, inferring their physical orientation. We validate these associations across biological contexts and reveal how polarization vector fields can be harnessed to discover multi-cellular structures (e.g. “streams” of cells with similar polarizations). These multi-cellular cytoskeleton dynamics associate with distinct phenomena, such as vector “divergence” with CD8^+^ T cell activation and “streams” with spatially segmented stages of the anti-tumor immune response. Together, these data support a role for CytoRipple in revealing a dynamic layer across spatial multi-omic platforms.

## Introduction

Spatial multi-omics have brought an unprecedented lens to the fields of biology^1^. Through the simultaneous measurement of a cell or molecule’s identity and spatial position, one can now directly interrogate for cellular or molecular interactions rather than infer their presence through dissociated data (e.g. single cell). Spatial multi-omic technologies vary in their plex (e.g. number of measured genes or proteins) and resolution (e.g. subcellular versus multi-cellular). While higher plex technologies have allowed for the spatially-resolved quantification of entire molecular pathways, this has often only been possible with lower resolution measurements (e.g. multi-cellular or >1 µm). While certainly a leap forward for the field, the limitations of these technologies hinder investigations into subcellular architecture which has recently been reported to play pivotal roles in cell biology^2–6^. Notably, an emerging set of spatial multi-omic technologies are seeking to address these limitations with spatially-resolved data that is both high in resolution and plex (e.g. Atera by 10x Genomics, whole-transcriptome CosMx by NanoString).

The links between subcellular heterogeneity and biological function have been well documented^7–10^. For example, autoimmune diseases can be distinguished by anti-nuclear antibody staining patterns^11^ and differences in nuclear architecture have been mapped to distinct cell states^4,5,12–14^. Cell movement drives some of the greatest changes in subcellular architecture as large-scale rearrangements of cytoskeleton elements are required to facilitate motility^15–17^. These changes manifest as cell polarization^18^, with preferential abundance of cytoskeleton proteins (e.g. actin, intermediate filaments) along the leading or trailing edge of the migrating cell^19^. Traditionally, subcellular protein polarization (hereby also called cytoskeleton dynamics) has been recorded through high-resolution, low-throughput imaging techniques^20–22^. The advent of spatial proteomics (e.g. Xenium, CosMx, Akoya PhenoCycler [CODEX], multiplexed ion beam imaging [MIBI]) has enabled high-throughput measurement of these cytoskeleton states within cells across whole tissues. In this work, we introduce a new computational framework, CytoRipple, that leverages these technological advances to generate tissue-scale maps of cytoskeleton dynamics. Through these maps, we uncover previously encoded relationships between cytoskeleton dynamics and biological phenomenon (e.g. limb development and immune cell infiltration of solid tumors).

## Results

### Overview of the CytoRipple Computational Framework

CytoRipple is a computational framework that utilizes subcellular measurements of cytoskeleton proteins to determine a cell’s polarization; this polarization often occurs in preparation for or during cell movement^15–19^ (**Fig. 1A, Methods**). CytoRipple quantifies cell polarization as a mathematical vector (with X- and Y-components) pointing the direction towards which the protein is most highly expressed cell; hereby, we term the angle of this vector as the polarization angle. We validated CytoRipple’s quantification of cytoskeleton polarization, and thus cell movement, using a published experimental model^23^ of immune cell (neutrophil) chemotaxis stained for actin, a cytoskeleton protein polarized at the anterior of the cell during migration. Using this model, we confirmed that the angle of cytoskeleton (actin) polarization over time, quantified by CytoRipple, was similar to the direction towards which the assayed neutrophil moved (**Fig. S1**).

**Fig. 1:**
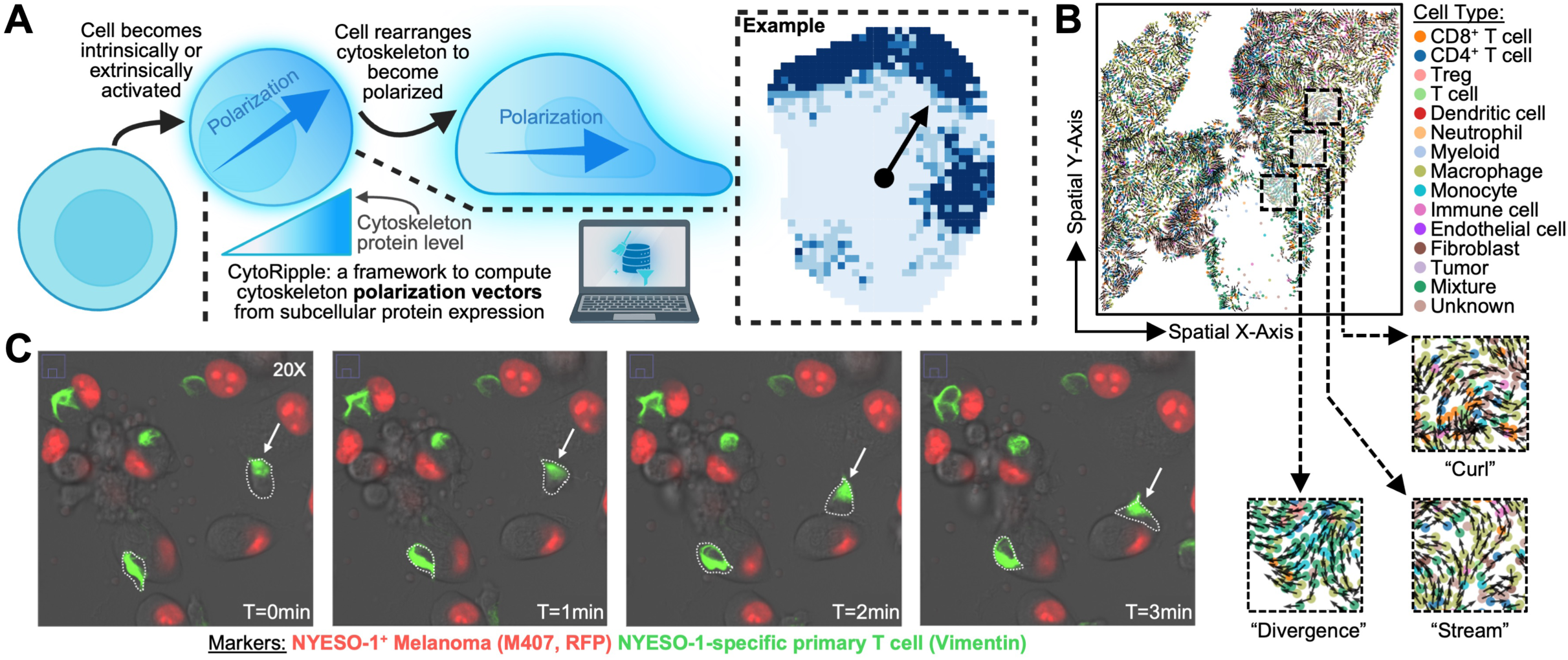
Overview of the CytoRipple Computational Framework. A) Diagram of the CytoRipple computational framework. Subcellular resolution measurements of cytoskeleton proteins (e.g. vimentin, tubulin, actin) were utilized to compute single cell polarization (i.e. the side of the cell cytoskeleton proteins are preferentially biased towards). B) Representative biopsy of polarization vectors (cytoskeleton dynamics) from a patient with melanoma treated with immunotherapy. Zoomed-in regions of the biopsy with representative multi-cellular cytoskeleton dynamics are outlined in dashed blue lines: streams of head-to-tail polarization vectors and regions of polarization vectors that have “divergent” directions. are presented on the right (dashed blue outline). Dots represent individual cells and arrows represent polarization vectors. Cells are colored by their assigned cell type (see legend on the right). C) Immunofluorescent imaging (20x) of vimentin stained (GFP [green]) NYESO-1-specific primary T cells with NYESO-1^+^ melanoma cells (M407 cell line, RFP [red]) at multiple timepoints of co-culture.

When applied at tissue-scale, using biopsies from a previously published cohort of patients with melanoma^24^, CytoRipple revealed cell polarizations to form distinct multi-cellular patterns that visually take on the appearance of “streams”, “curls”, and “divergences” (**Fig. 1B**). “Streams” characterize cells that are polarized in the same direction and form a head-to-tail line. “Curls” and “divergences” are multi-cellular patterns that exhibit their namesake behavior and can be quantified mathematically (**Methods**). Given the focus of our downstream analyses on cancer immunotherapy contexts, we sought to validate the existence of cell polarization in involved cell types. To this end, we stained for the cytoskeleton protein vimentin in T cells engineered to express the T cell receptor (TCR) specific to NY-ESO-1 co-cultured with melanoma cells expressing NY-ESO-1 and indeed observed cell polarization by live cell imaging during migration and target tumor-cell engagement (**Fig. 1C**). While CytoRipple can be applied to any protein, vimentin was the primary cytoskeleton protein used for downstream analyses due to the ubiquity of this protein in panels across technologies (including 10X Xenium, multiplexed immunofluorescence, CODEX, and MIBI in this study) and its known role in motility-associated cell polarization^25,26^.

### Validation of Model Design and Parameter Choice

Prior to applying CytoRipple to biological use cases, we first sought to confirm the model’s computational approach. To this end, we leveraged our prior dataset of published biopsies from patients with melanoma^24^ to characterize CytoRipple’s mathematical quantification of multi-cellular patterns (“curl” and “divergence”) and parameter choices. To compute “curl” and “divergence”, an NxN grid is overlaid on a tissue of interest and said values are computed for each block within this grid (**Fig. S2A**). Computed values were then interpolated to create high-resolution maps of “curl” and “divergence” (**Fig. 2A**). Grids large enough to encompass multiple cells are required to have enough data points to perform these computations. As grid size is a tunable parameter, we investigated how “curl” and “divergence” values varied with changes in grid size (e.g. 10x10 versus 20x20 grid). No substantial differences in computed values for “curl” and “divergence” were observed when different grid sizes were utilized for said calculations, average Z-score difference of 0.36±0.07 (mean ± 95% confidence interval) and 0.36±0.07 for “curl” and “divergence”, respectively (**Fig. 2B**). A grid size of 100x100 was utilized for subsequent analyses to maximize resolution, maintaining a sufficient number of cells per grid block to perform “curl” and “divergence” calculations.

**Fig. 2:**
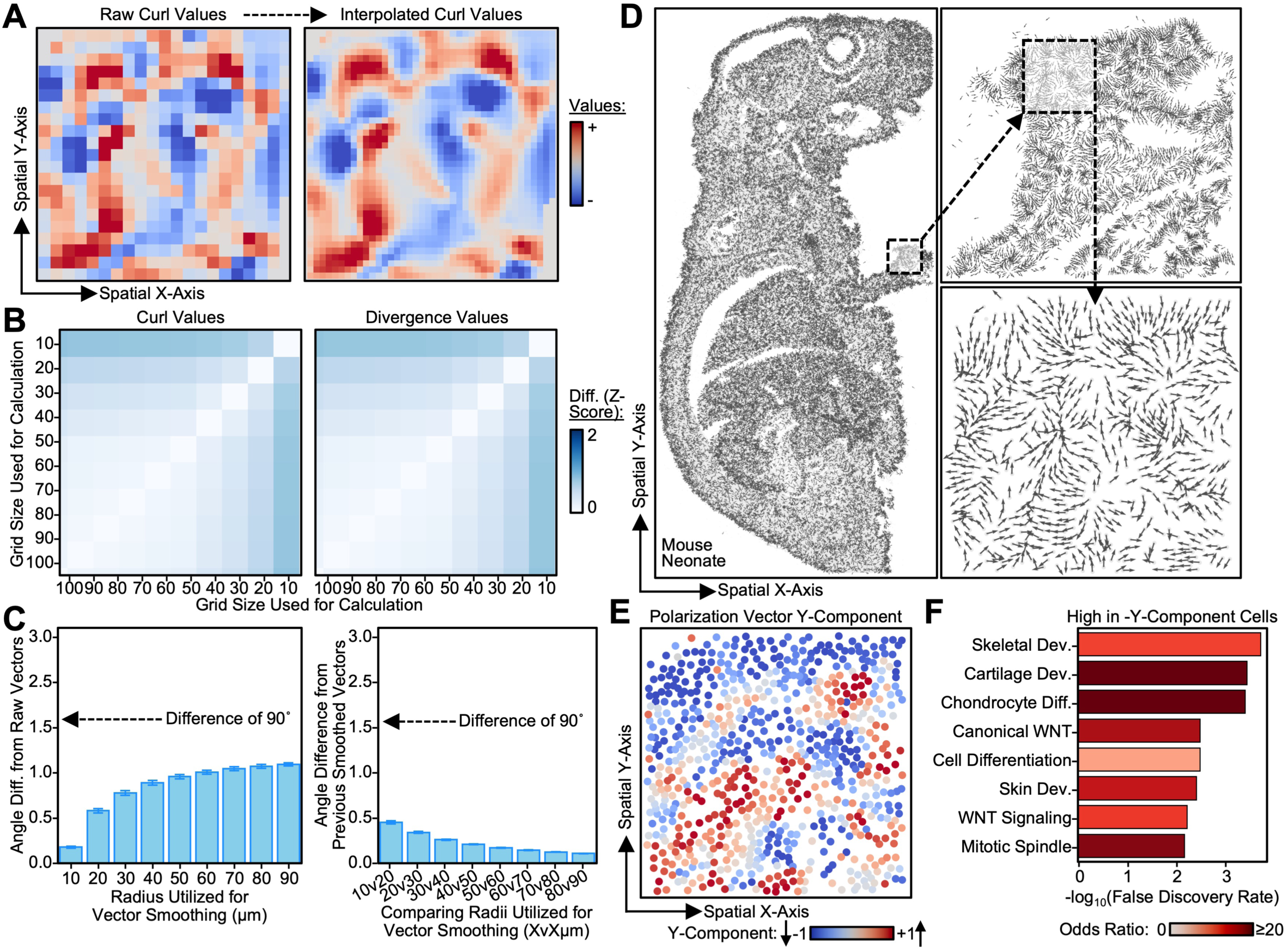
Technical and Biological Validation of CytoRipple Parameters and Model Design. A) Spatial plot of raw (left) and interpolated (right) curl values (see legend on the right). B) Heatmap of the difference in curl and divergence values computed using varying grid sizes (see legend on the right). Curl and divergence values were Z-score normalized to allow for fair comparisons between grid sizes. C) Bar plots of the differences in pre- and post-smoothing polarization angles (in radians, Y-axis) using a range of radii to perform said smoothing (X-axis). The difference between pre- and post-smoothing angles on the left. Relative differences between post-smoothing angles are presented on the right (e.g. at X=20µm, the Y-axis represents the difference between the post-smoothing polarization angles computed using a radius of 10µm versus 20µm). D) Keft: Spatial plot of whole mouse pup. Dots represent individual cells and arrows represent their polarization vectors. Vectors were downsampled (to N=50,000) for visualization purposes. Right: Zoomed-in sections of the developing limb and region of interest (ROI). This ROI was chosen due to growth processes reported to occur at the edges of developing limbs. E) Spatial plot of the Y-component of the polarization angles from single cells in the developing mouse limb’s region of interest from Fig. 2D (see legend on the bottom). F) Bar plot of pathways (Y-axis) enriched in cells polarized to move downwards (negative Y-components). Enrichment is quantified by -log_10_ false discovery rate (FDR, X-axis). Color represents odds ratio (see legend on the bottom). Bar heights represent arithmetic mean and error bars represent 95% confidence interval.

CytoRipple performs spatially-weighted smoothing of polarization vectors (**Fig. S2B**). Smoothed polarization vectors are computed by normalizing each original vector by the average of its neighboring vectors within a fixed spatial radius. This smoothing reduces technical noise to maximize biological signal, similar to previous computational frameworks applied to other technologies^27,28^. The fixed radius is a tunable parameter. To identify an ideal radius for vector smoothing, we investigated how pre- and post-smoothing angle differences varied by radius size (**Fig. 2C**). We noted a sharp increase in pre- and post-smoothing angle differences at a radius of 20 µm, likely corresponding to the minimum cell radius (greater than the size of an average cell)^29^ that captures nearby cells. At radii greater than 20 µm, pre- and post-smoothing angle differences exhibited minimal changes; this supports 20 µm as the minimum distance to capture vectors from a non-negligible number of spatially proximal cells. Notably, none of the radii led to pre- and post-smoothing angle differences to be altered by greater than 90° on average. To balance preserving the angles of the original polarization vectors while reducing technical noise through smoothing, all downstream analyses were performed with a radius of 30 µm.

### Validation of CytoRipple through Biological Case Studies

To validate CytoRipple, we applied this framework to two biological contexts where the role of vimentin has been previously described: mouse limb development^30^ and colonocytes in human intestine^31^ (**Data Availability**). For the case study on the developing limb, we applied CytoRipple to a Xenium dataset of a whole mouse pup (one day old) and focused in on a section of the growing paw (**Figs. 2D and S2C**). We found developing limb cells to largely group into those polarized distally (away from the body) and proximally (**Figs. 2E and S2D**). Consistent with its known role in limb development, WNT signaling was enriched in proximally polarized cells (cytoskeletons oriented towards the body, **Fig. 2F and Tables S1-S2**). For the case study on colonocytes, we applied CytoRipple to a CODEX dataset of healthy human intestine^31^. We observed colonocytes to present with basally polarized cytoskeletons, consistent with independent protein staining (immunohistochemistry, **Fig. S3**). Together, these data suggest that the cytoskeleton dynamics identified by CytoRipple recapitulate known biological phenomena.

### Cytoskeleton Dynamics Associate with Distinct Molecular Phenotypes in Melanoma

Having validated CytoRipple’s ability to recapitulate known biology, we next interrogated if CytoRipple could identify new biology phenomenon linked to cytoskeleton dynamics. To achieve this, we applied CytoRipple to our previously published cohort of melanoma biopsies from patients before and during treatment with immunotherapy (n=220 regions of interest)^24^. We observed marked heterogeneity in cytoskeleton dynamics across tissue biopsies; these dynamics seemed to visually correspond with distinct cell types (**Fig. 3A**). To formally investigate this association, we correlated metrics of cytoskeleton dynamics (e.g. “curl”) with the location-matched protein expression within each biopsy, across all 32 proteins stained (**Figs. 3B and S4A**). As expected, expression of the cytoskeleton protein vimentin strongly correlated with the magnitude of its polarization vector. Interestingly, we also observed certain immune-related proteins associating with distinct multi-cellular patterns. For example, we found B cell marker CD20 and neutrophil marker MPO to correlate with “convergence” (negative “divergence”) across biopsies.

**Fig. 3:**
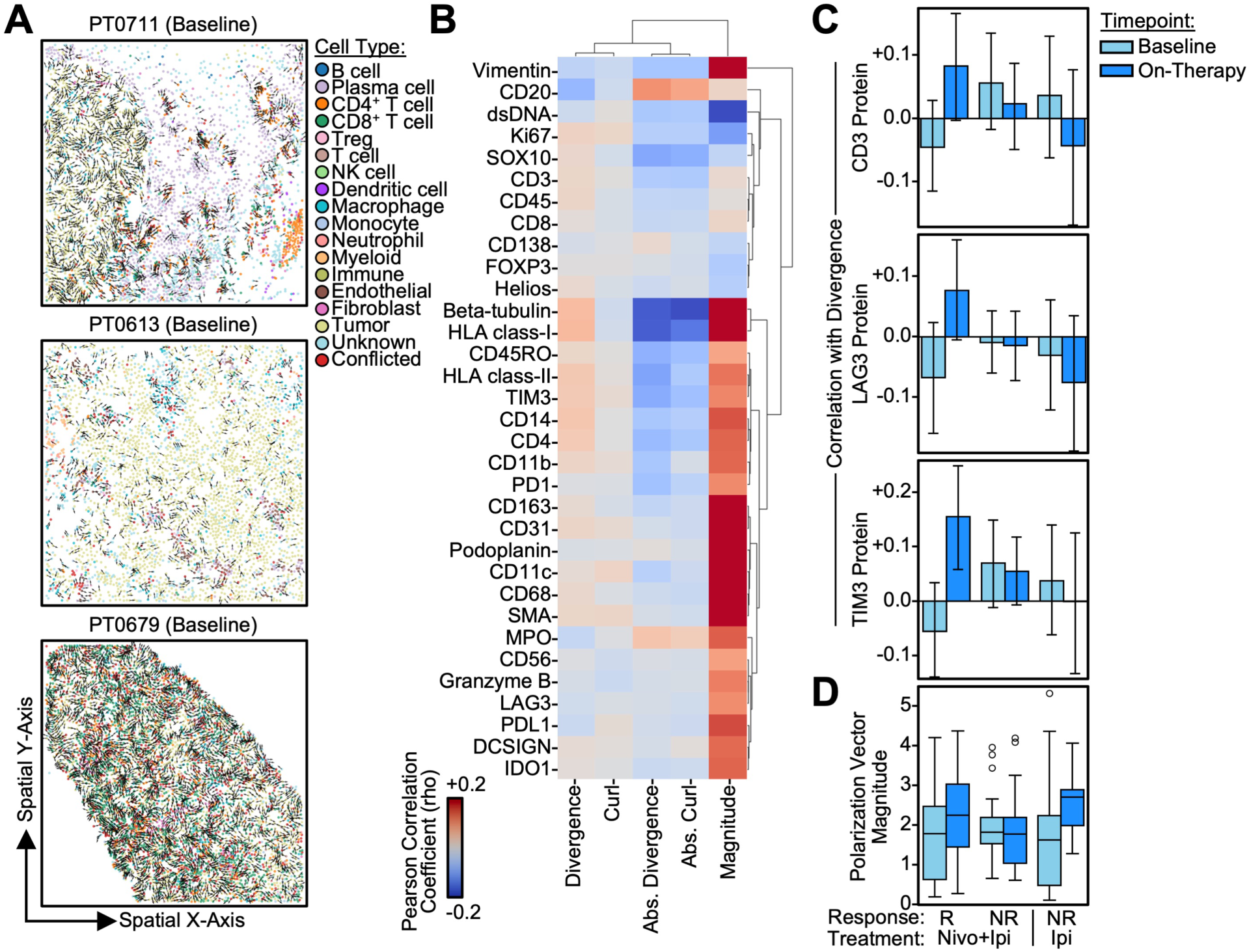
Cytoskeleton Dynamics Associate with Heterogeneous Protein Expression Patterns in Human Melanoma. A) Spatial plots of representative biopsies from patients with melanoma. Each dot represents a single cell, and arrows represent their polarization vectors. Dots are colored by their assigned cell type (see legend on the right). B) Heatmap of the correlation (Pearson’s method) between protein abundance (rows) and cytoskeleton dynamics (columns) in patients with melanoma treated with immunotherapy. Color represents the average correlation coefficient (rho) across samples (see legend on the bottom left). C) Bar plots of the coefficients of correlation (rho) between divergence (a cytoskeleton dynamic) and protein expression (Y-axis) as stratified by patient treatment and response (X-axis). Color represents timepoint: pre-treatment in dark blue and post-treatment in light blue (see legend on the left). Bar heights represent arithmetic mean and error bars represent 95% confidence interval. D) Box plot of polarization vector magnitude (i.e. the magnitude of vimentin protein expression in the direction of the vector, Y-axis) as stratified by patient treatment and response (X-axis). Color represents timepoint: pre-treatment in dark blue and post-treatment in light blue (see legend on the left). Each dot represents an individual sample. Box plot represents interquartile range (25th to 75th percentiles, IQR) with whiskers representing 1.5 times IQR.

Given that immune cell infiltration and multi-cellular neighborhoods vary based upon immunotherapeutic response, we next queried whether the associations between cytoskeletal dynamics and protein expression varied across patient responses to immunotherapy. For example, do CD8 protein levels only associate with increased “convergence” in certain patient groups? Our analysis revealed biopsies from patients who responded to combination immunotherapy (anti-PD1 and anti-CTLA4) to have increased correlation between cell “divergence” and expression of T cell (CD3) and T cell exhaustion associated (LAG3, TIM3) proteins (**Figs. 3C and S4B**). This may correspond to the known stop-and-go (i.e. “divergent”) motility pattern of tumor-specific T cells^32^ which are also known to upregulate exhaustion markers after chronic antigen stimulation^33,34^. We verified that our findings were not simply due to differences in cytoskeleton protein expression (i.e. polarization vector “magnitude”, **Fig. 3D**). In all, global and cell type specific cytoskeleton dynamics bear distinct associations with cellular proteomic profiles and patient responses to immunotherapy.

### Multi-Cellular Polarization Patterns Identify Immunological Niches in Melanoma

Cells do not exist in isolation; their spatial surroundings influence their fate and function (e.g. via secreted cytokines)^35^. Thus, we investigated if these single-cell cytoskeleton dynamic associations reflected broader regional shifts in tissue biology. To achieve this, we subjected each tissue biopsy to spatial and cytoskeleton dynamic informed clustering, based upon the position and directionality of the vimentin vector fields (**Methods**). Unbiased clustering revealed 9,323 identified regions across 220 images, which grouped into 16 Leiden clusters based upon cytoskeleton dynamics and proteomic profiles (**Figs. 4A-B**). Interestingly, one regional cluster (0) simultaneously presented with polarization vector “divergence”, expression of CD8, cytotoxic granzymes (GZMB), T cell exhaustion related proteins (PD1, LAG3, and TIM3), and IFNG-induced proteins (HLA class II and PD-L1). Cell type analysis of regional cluster 0 confirmed the cluster’s significant enrichment of CD8^+^ T cells (**Fig. 4C**). Together, these data suggest this region as representative of highly-motile (“divergent”) or activated, antigen-experienced (PD1, and TIM3 expression) cytotoxic (GZMB) CD8^+^ T cells located in an IFNG-rich spatial environment (HLA class II, **Fig. 4D**). Consistent with their seemingly pro-tumoricidal biology, regional cluster 0 was significantly more prevalent in baseline biopsies from patients who responded to immunotherapy (**Fig. S5**). In all, we demonstrate how CytoRipple synergizes with spatial analysis frameworks (e.g. regional clustering) to identify immunotherapy response-associated spatial niches and provide cytoskeleton dynamic informed characterization of these niches, such as “divergent” polarization vectors suggestive of motility or activation.

**Fig. 4:**
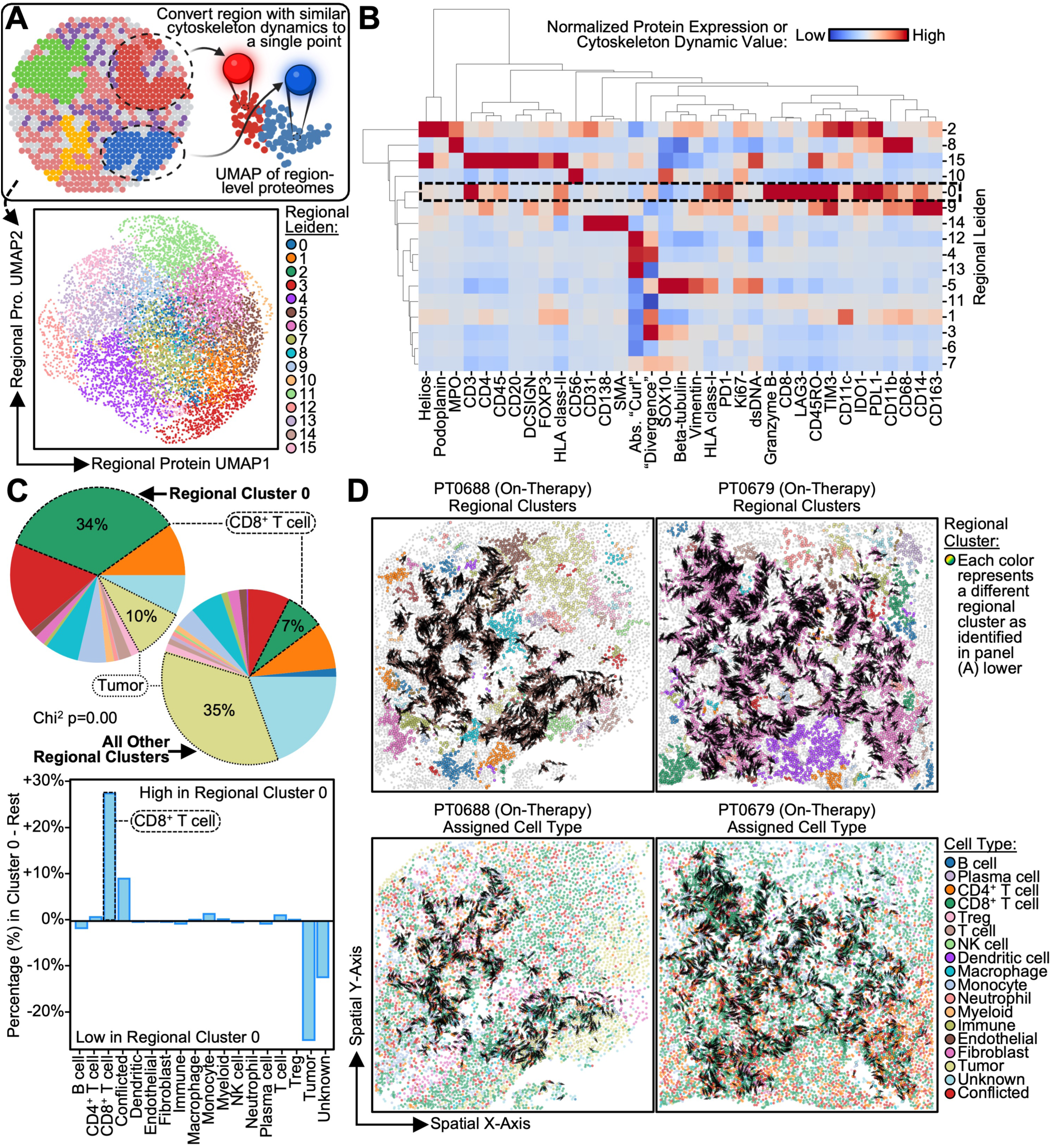
Regional Cytoskeleton Dynamics Identify an Immunologically “Hot” Spatial Niche Characterized by Divergent Cell Polarizations. A) Upper: Diagram demonstrating how single cells were clustered into spatial regions by their cytoskeleton dynamics and protein expression. Lower: Uniform manifold approximation projection (UMAP) of regional cytoskeleton dynamic and protein expression profiles. Each dot represents a single region from an individual sample. Color represents unbiased clustering of these regions (see legend on the right). B) Heatmap quantifying the degree of protein expression and cytoskeleton dynamics (columns) of each unbiased cluster of regions from panel (A) (rows). Color represents protein expression or cytoskeleton dynamic magnitude (see legend on the right). Dashed black outline indicates a regional cluster of interest (cluster 0) that is simultaneously enriched for CD3 and CD8 proteins and divergence of polarization vectors. C) Upper: Pie plots of the distribution of cell types in single cells that are in regional cluster 0 (left) compared to those that are in all other regional clusters (right). Color indicates assigned cell type (see legend on the bottom). Bottom: Bar plot of the difference in distribution (percentage [%]) of cell types in regional cluster 0 (positive values) compared to cells in all other regional clusters (negative values). P-value was calculated using the Chi-squared test. D) Spatial plots of representative biopsies with regions classified as regional cluster 0; cells in this cluster are outlined in black. Arrows represent polarization vectors from the cells in regional cluster 0. Cells are colored by their regional cluster on the top and assigned cell type on the bottom (see legends on the right).

### Cytoskeleton Dynamics Reveal Heterogeneous Cellular Trajectories in Melanoma

Multi-cellular cytoskeleton dynamics can not only be observed as regional shifts but also as “streams” (**Fig. 1B**). That is, cells aligned head-to-tail that bear similar cytoskeleton polarizations. We hypothesized that cells oriented as localized streams may be responding to a common stimulus (e.g. chemokine)^36^. To assign cells to streams, a cell is chosen (“origin”) and progressive sets of cells are included if there exist cells that both lie in the direction of the chosen cell’s polarization vector and bear a similar polarization angle (45° in either direction, **Fig. 5A**). To better understand the molecular features associated with this behavior, we systematically computed these streams, if present, for each cell in a representative on-treatment biopsy with 34% tumor cellularity (area positive for melanoma marker S100) from a patient with melanoma responding to combination immunotherapy (PT0242, on-therapy, **Methods**). We detected 1,922 streams (compared to 16.8±6.9 by a null model) that originated, traversed, and terminated on a convergent set of locations in the tumor (**Figs. 5B-C**). When we interrogated for commonalities between these streams, assigning the intratumoral endpoint of each stream as the “origin,” we found that the termini across streams generally had the highest level of immune-related proteins (**Fig. 5D and S6A**). When streams were compared based upon their normalized protein expression, they were grouped into 20 Leiden clusters, revealing subtle differences in immune streams within the tumor microenvironment (**Figs. 5E and S6B**). Leiden 7 was Treg dominant, with high FOXP3 and Helios expression at the termini and Ki67 and CD45RO concentrated within the stream and towards the termini (**Fig. 5F**). Alternatively, Leiden 8 was myeloid dominant, with high expression of CD11b, CD68, CD14, and CD4 throughout the streams (**Fig. 5F**). These differential termini may represent different stages of tumor resolution, such as ongoing tumoricidal activity in T cell enriched termini or wound healing following tumor cell killing in termini with myeloid predominance^37,38^. These analyses demonstrate how CytoRipple may be utilized to identify heterogeneous cellular trajectories within tissue biopsies.

**Fig. 5:**
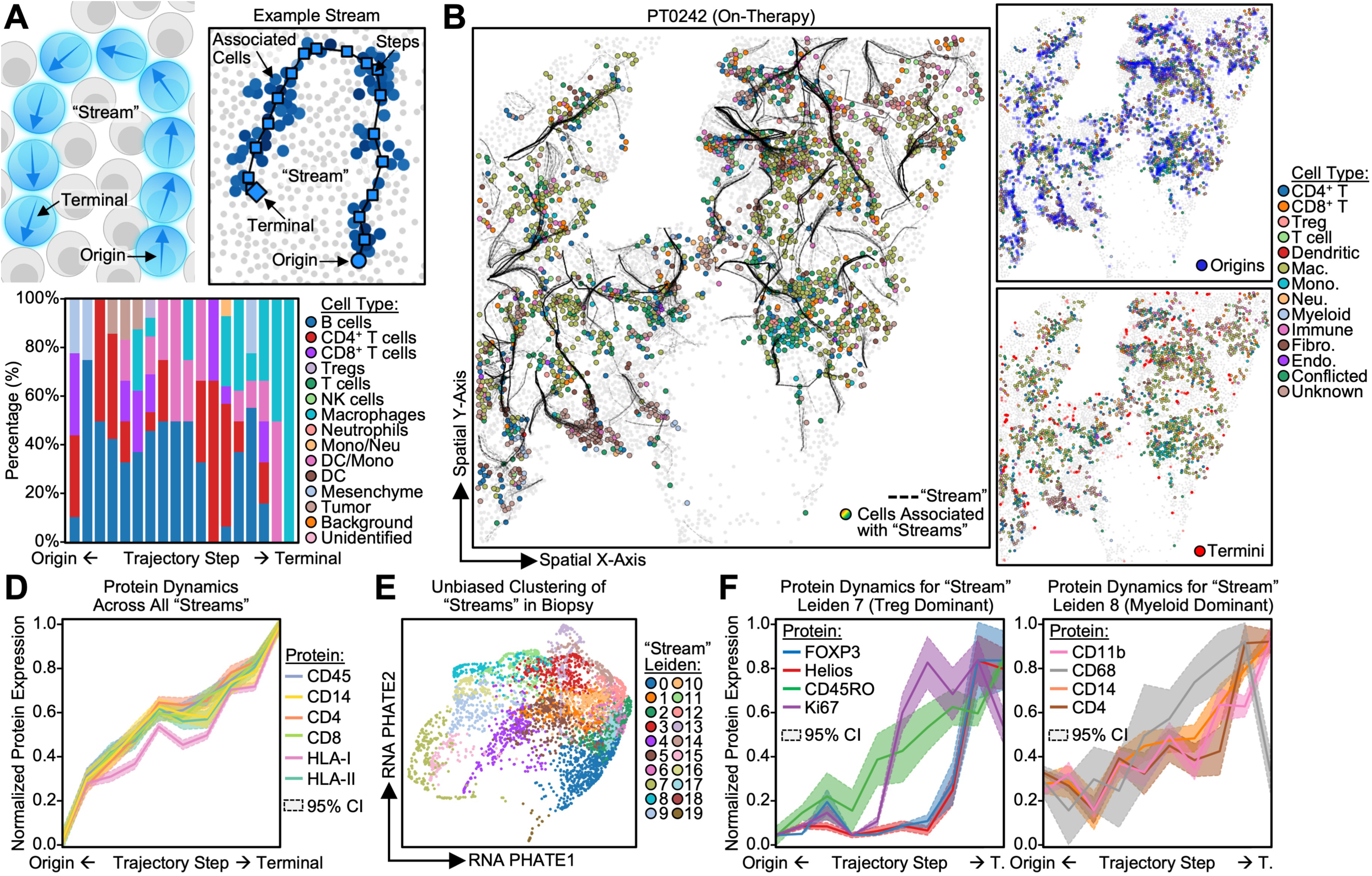
Multi-Cellular Cytoskeleton Dynamics Reveal Heterogeneous Spatial Trajectories in Human Melanoma. A) Diagram of the computational identification of spatial trajectories (streams) from multi-cellular cytoskeleton dynamics (upper left). A representative stream highlighting the origin (circle), intermediate steps (square), and terminal (diamond) with the associated cells at each step colored in blue; cells further away from the stream are colored in lighter blue (upper right). Bar plot of the distribution of cell types at each step along the trajectory of the representative stream (bottom); color represents assigned cell type (see legend on the right). B) Spatial plot of the streams (dashed black lines) in a melanoma biopsy with cells associated with each stream colored by their assigned cell type (see legend on the right of panel [C]). Overlapping streams appear as solid black lines due to the overlap of many dashed black lines. C) Spatial plots of the origins (upper) and termini (lower) of each stream in the melanoma biopsy presented in panel (B). Cells associated with each stream are colored by their assigned cell type (see legend on the right). D) Line plot of the average protein expression (Y-axis) at each trajectory step across all streams (X-axis) in the melanoma biopsy presented in panel (B). Line color represents protein identity (see legend on the right). Shaded area represents 95% confidence interval. E) UMAP of trajectory steps across all streams in the melanoma biopsy presented in panel (B). Dots represent trajectory steps that are numerically represented by the protein expression of nearby cells and the protein expression of cells proximal to the trajectory steps before and after the given trajectory step. Dots are colored by their unbiased trajectory cluster assignment (see legend on the right). F) Line plots of average protein expression (Y-axis) at each trajectory step across streams (X-axis) in trajectory cluster 7 on the left and trajectory cluster 8 on the right. Line color represents protein identity (see legend on the right). Shaded area represents 95% confidence interval.

### Immune Cell Trajectories Associate with Chemoattractant Gene Expression in Lung Adenocarcinoma

To validate CytoRipple’s ability to identify biologically meaningful cellular trajectories (streams), we applied this framework to a public, spatially-resolved single-cell transcriptomic dataset (10X Xenium platform) of human lung adenocarcinoma (**Figs. 6A and S7A, Data Availability**). We applied CytoRipple to the multiplexed immunofluorescence image corresponding to vimentin staining from the 10X Xenium experiment performed on this biopsy and identified 2,758 streams (compared to 1.0±0.7 by a null model, **Figs. 6B-C**). Streams were enriched for the presence of T cells and expression of immune cell stimulating and recruiting chemokines (e.g. *CCL5*^39^ and *CXCL9*^40^, **Figs. 6D and Table S3**), while cells outside of streams were enriched for tumor cells and corresponding expression of *EPCAM, KRT7,* and *EGFR.* Streams were clustered based upon the gene expression along the cells in their trajectory (**Fig. 6E**). This revealed homogenous mapping of streams with origins in tumor regions (*EPCAM^+^*) and termini reflecting expressing genes from immune cells and vasculature (*CD3E*^41^ and *PECAM1*^42^ **Fig. 6F**). Interestingly, the cells at the origin of the stream also demonstrated expression of immunosuppressive chemokines (e.g. *CXCL2*^43^, **Fig. 6F**).

**Fig. 6:**
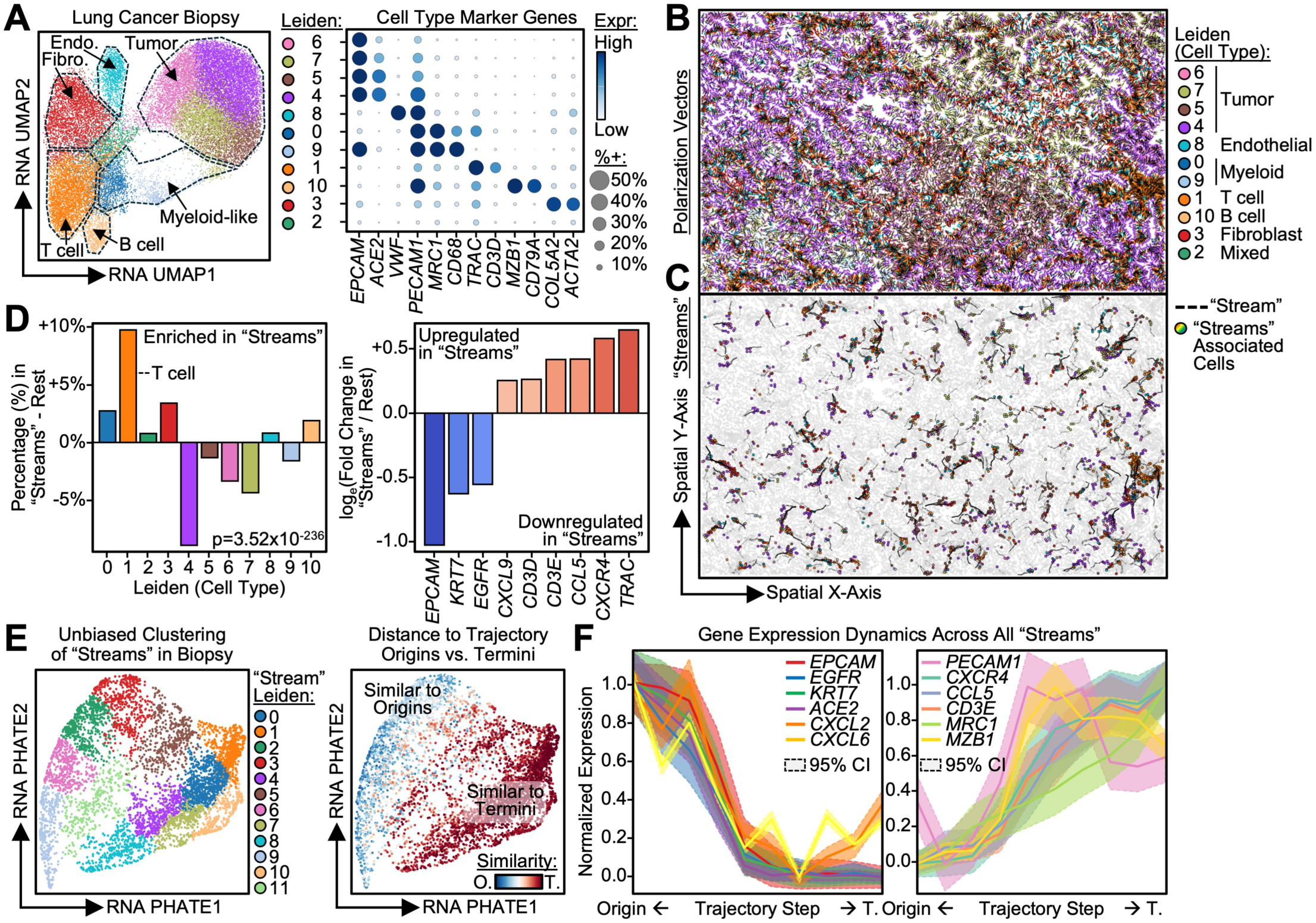
Validation of the Presence of Spatial Trajectories “in Human Lung Adenocarcinoma. A) Left: UMAP of single cell transcriptomes from a biopsy of human lung adenocarcinoma. Dots are colored by their unbiased (leiden) clustering (see legend on the right) with the cluster’s assigned cell type annotated directly on the plot (dashed outlines). Right: Dot plot of the marker genes (columns) used to classify unbiased clusters (rows). Dot size indicates the fraction of cells in a given cell type expressing a given gene and dot color represents average expression (see legend on the right). B) Spatial plot of the biopsy in panel (A). Each dot represents a single cell, and arrows represent their polarization vectors. Dots are colored by their unbiased cluster (see legend on the right). C) Spatial plot of the streams (dashed black lines) from the biopsy in panel (A) with cells associated with each stream colored by their unbiased cluster (see legend on the right). Overlapping streams appear as solid black lines due to the overlap of many dashed black lines. D) Left: Bar plot of the difference in distribution (percentage [%]) of unbiased clusters in streams (positive values on the Y-axis) compared to cells not in streams (negative values on the Y-axis). P-value was calculated using the Chi-squared test. Right: Bar plot of the differentially expressed genes (X-axis) in cells associated with streams. Y-axis quantifies differential expression as log_2_ fold change (FC) of the average expression in “stream” associated cells over the average expression in all other cells. Upregulated genes have positive values on the Y-axis (red, color corresponds to log_2_FC) while downregulated genes have negative values (blue). All presented genes are significant (False Discovery Rate [FDR] < 0.05). P-values were calculated using the Mann-Whitney U test. E) Left: UMAP of trajectory steps across all streams in the biopsy presented in panel (C). Dots represent trajectory steps that are numerically represented by the gene expression of nearby cells and cells proximal to the trajectory steps before and after the given trajectory step. Dots are colored by their unbiased trajectory cluster assignment (see legend on the right). Right: UMAP of trajectory steps colored by their expression similarity (quantified by Euclidean distance) to cells that are origins of streams (similar cells are blue) vs. cells that are similar to termini of streams (similar cells are red, see legend on the bottom right). F) Line plot of the average gene expression (Y-axis) at each trajectory step across all streams (X-axis) in the biopsy presented in panel (C). Line color represents transcript identity (see legends in each plot of the panel). Shaded area represents 95% confidence interval.

The anti-tumor immune response is reported to occur in spatially sequential stages; immune cells extravasate from vasculature prior to interacting with tumor cells, with T cells trafficking further into the tumor^44–47^. CytoRipple streams qualitatively align with these known stages via enrichment of vasculature and tumor markers at opposite ends of identified trajectories (**Fig. 6E**). This prompted us to investigate this similarity in a more quantitative fashion. To investigate this, we identified co-expressed gene modules across each “stream” (**Figs. S7B-C, Tables S4-S5**) mapping their positions along each cellular trajectory (**Fig. S7D**). This revealed a picture similar to the reported spatial stages of the anti-tumor immune response. One end of the trajectory was enriched for tumor cell genes (modules 1 and 6), while the other terminus was enriched for vasculature genes (module 2), T cell genes (module 4), and other immune cell genes (module 5). Collectively, this is suggestive of the orientation of the cells associated with inflammatory recruitment due to anti-tumor immunity. In all, this data suggests that CytoRipple may identify cellular trajectories in tumor biopsies representative of dynamic processes at the tissue level.

## Discussion

Chemotaxis plays a significant role in facilitating the ever-shifting landscape of immune, stromal, and tumor cell interactions that define the tumor microenvironment^38,48,49^. These interactions can both promote tumor cell death (e.g. via recruitment of anti-tumor T cells) and tumor cell growth (e.g. through the attraction and retention of immunosuppressive neutrophils)^50^. Historically, these chemotactic processes have been observed through high-resolution microscopy techniques^20,51^ capable of imaging the cytoskeleton polarizations that facilitate cell movement towards or away from chemotactic stimuli^15–19^. In this work, we reveal how existing cytoskeleton protein measurements in spatial-omic assays can be leveraged to uncover previously hidden tissue-scale maps of these cytoskeleton dynamics, and thus spatially map chemotactic processes^52^. By quantifying cell polarizations from multi-omic data (e.g. Xenium, CODEX, and MIBI), CytoRipple facilitates deep molecular interrogations of chemotactic processes; these investigations were previously limited by the handful of measured markers and field of view compatible with traditional microscopy techniques. In addition, we quantify the well-discussed concept of cellular streams: coordinated groups of cells with similar cytoskeleton polarizations that are likely responding to shared chemotactic stimuli^53,54^.

Through a series of case studies, we revealed how this new view of chemotactic processes allows for confirmation of previously known biology (e.g. enrichment of CD8^+^ T cells in streams) and discovery of new phenomenon (streams of Treg- versus myeloid-rich termini). Interestingly, when we quantified cellular responses along tumor biopsy streams, we identified a spatially sequential set of processes reminiscent of the known stages of the anti-tumor immune response^44–47^: early vasculature, subsequent myeloid cell enrichment, T cells at the tumor border, and then the tumor cells themselves. It has been hypothesized that tumors resist clearance by the immune system through disruption of these stages. For example, although fibroblasts can limit metastatic spread by creating a fibrous matrix around the tumor, they can also limit tumor resolution by creating a physical barrier between tumor cells and anti-tumor cytotoxic T cells^55^. Similarly, macrophages and neutrophils aid in wound healing after cancer cell clearance when on the tumor periphery, but tolerance-polarized myeloid cells can suppress tumoricidal T cells when localized to the tumor core^56–59^. We anticipate that these stages of anti-tumor immunity will be increasingly well understood by evaluating not just cell types and states, but the dynamic inference defined by their morphology and cytoskeletal orientation.

### Limitations of the Study

This study is limited by its reliance on proper segmentation. Polarized cells often have atypical shapes that can be difficult for typical computational frameworks to segment properly, either by improved cell membrane markers or higher resolution imaging. Despite this challenge, we do see recurrent patterns (i.e. streams, curl, divergence) across Xenium, CODEX, and MIBI datasets, each with unique panels, resolution, and segmentation algorithms. Further, certain technologies measure multiple proteins in the same fluorescent channel (e.g. vimentin/SMA for cell segmentation by Xenium). While we did not find this to majorly affect our findings in this work, future studies where SMA plays greater biological roles may need to consider alternate cytoskeleton proteins or measurement modalities. Additionally, although the direction a cell is moving often aligns with its angle of polarization, there are certain cell type specific exceptions that may need to be considered for future works that are explicitly mapping cell motility rather than the tissue-scale trajectories we discuss in this work. Finally, while cytoskeleton polarization is required for cell motility, it does not always indicate a motile cell. For example, certain cytoskeleton proteins serve as physical anchors to keep cells in their proper physiological place (e.g. hemidesmosomes in the skin).

## Methods

### CytoRipple Computational Framework

The complete CytoRipple package is provided freely for public usage on GitHub at: https://github.com/danielgchen/CytoRipple with scripts required to reproduce all analyses presented in this manuscript at: https://github.com/danielgchen/reproducibility_cytoripple. Here, we provide an in-depth description of the CytoRipple computational framework to accompany the aforementioned code. The framework follows the following pseudo-code: 1) Read in the cytoskeleton protein of interest (e.g. actin, vimentin), 2) Raw polarization vectors are computed using each cell’s subcellular distribution of the chosen cytoskeleton protein, 3) Polarization vectors are subject to spatial smoothing (further description below), 4) Smoothed polarization vectors can be subject to “curl” and “divergence” calculations to identify multi-cellular cytoskeleton dynamics (further description below). X- (or Y-) component of raw polarization vectors are computed as the difference in subcellular cytoskeleton vector expression between right and left (or lower and upper) halves of the cell of interest; coordinates of the halves are derived from the centroid of the cell.

### *In Vitro* Validation of T Cell Cytoskeleton Polarization in Tumor Cell Co-Culture

Patient-derived melanoma cells expressing NY-ESO-1 (M407 cell line^60^) and stably transduced with a nuclear RFP expression vector (M407-nRFP cells) described previously^61^ were cultured in DMEM (Corning #10-013-CM) with 10% FBS (Cytiva HyClone #SH30071.03) at 37°C and 5% CO2.

Primary CD8+ T cells were isolated, activated, and transduced with an HLA-matched NY-ESO-1-specific transgenic T cell receptor (TCR) as previously described2. Briefly, peripheral blood mononuclear cells (PBMCs) were thawed, and CD8+ T cells were isolated using a CD8+ T Cell Isolation Kit (Miltenyi Biotec #130-045-101). These T cells were then activated with Dynabeads Human T Activator CD3/CD28 (Invitrogen #11132D) and 50IU/mL human IL-2 (Miltenyi Biotec #130-097-748). Activated T cells were transduced with an NY-ESO-1-specific TCR retroviral expression vector in 6 well plates precoated with RetroNectin (Takara #T100B). They were additionally transduced with a commercially available vimentin-GFP overexpression lentiviral vector (Sigma-Aldrich #17-10152) at a high MoI. These T cells were spinoculated for 10 minutes at 2000xg. T cells were cultured and expanded in RPMI (Corning #10-040-CM) with 10% FBS and 50IU/mL hIL-2 at 37°C and 5% CO2.

The night prior to the experiment, M407-nRFP cells were split and plated at 10,000/250µL in 8 well chamber slides (Ibidi #80826). The following morning, the medium was replaced with fresh medium containing 10,000/250µL Vim-GFP CD8^+^ T cells. The coculture was incubated at 37°C for 1.5 hours prior to imaging. Imaging was performed with a Nikon AX R scanning confocal microscope with an incubator stage keeping the cells at 37°C and 5% CO2 for the duration of the experiment. 3 channel images (GFP, RFP, and brightfield) were taken every minute for 30 minutes at 20X magnification.

### CytoRipple Polarization Vector Processing

Regarding the spatial smoothing of polarization vectors, each cell’s polarization vector is recomputed as the weighted average of its original vector and the vectors of cells spatially proximal to the given cell of interest. A set distance is provided to the program, as a parameter, to determine the maximum distance spatially proximal cells can be from the cell of interest to be considered for the spatial smoothing. Spatial smoothing is performed in a weighted manner, such that the original cell’s polarization vector is given the greatest importance while cells at increasing distances away from the original cell are given smaller and smaller weights. This strategy was implemented as cells with decreased physical proximity to the original cell are less likely to be reacting to the same biological signal (e.g. chemokine) the original cell is reacting to (i.e. the cause of the original cell’s cytoskeleton polarization). Further, spatial smoothing was implemented as spatial multi-omic data is often noisy and thus the averaging of individual data points into broader trends allows for the emergence of robust signals and diminishment of true noise. For example, vectors pointing broadly in the same direction will have their signal amplified while vectors that point in a plethora of directions will have their signal reduced.

### CytoRipple Multi-Cellular Cytoskeleton Dynamics Computation

Regarding the computation of multi-cellular cytoskeleton dynamics, we computed for two main multi-cellular cytoskeleton dynamic features: “curl” (polarization vectors that appear to form a curl) and “divergence” (polarization vectors that point outwards or inwards from or to a common source, respectively). We address the computation of the multi-cellular structures of streams and “regions” in later sections of the Methods. To compute curl and divergence, a grid of cells is required; this was investigated in-depth in Fig. 2 whereby multiple grid sizes were tested to ensure that computed “curl” and “divergence” values did not significantly differ due to this technical parameter. Notably, it is worth considering increased grid sizes (e.g. 100x100 instead of 10x10) to increase resolution, however interpolation strategies seem to ameliorate this issue (i.e. as tested in this manuscript by comparing interpolated 10x10 grids to 100x100 grids). To mathematically compute for “curl” and “divergence”, we calculated the derivative of the X- and Y-component of each vector as a function of spatial X- and Y-dimension, respectively. “Curl” was defined as the difference between these two derivatives as a curl is defined by an increase in change in X with a concomitant decrease in change in Y (or vice versa). “Divergence” was defined as the sum of these two derivatives for increased changes in X and Y is the mathematical expression of vectors pointing away from a common point; negative “divergence” is thus equal to vectors pointing to a common spatial point (i.e. “convergence”). The creation of grids with different aspect ratios for spatial slides of different aspect ratios was not considered in this manuscript but may be easily changed through our freely available code by adjusting the “NGRIDX” and “NGRIDY” constants.

### Validation of CytoRipple in Multiple Contexts

Validation of CytoRipple using neutrophil actin polarization post chemokine stimulus was completed using publicly-available videos of such from *Weiner et al.*^23^. Each frame of the video, an image, was processed via FIJI^62^ to segment cells and export their spatial positions and subcellular cytoskeleton (actin) intensities. This data was then subject to the standard CytoRipple pipeline described in the prior Methods section. The angle of each polarization vector (hereby termed polarization angle) was computed by applying the tangent function to the ratio of the Y- and X-component of each polarization vector. Calculated angles were then normalized to 0-360 degrees (i.e. 0-2π radians).

Validation of CytoRipple using mouse pup vimentin polarization was completed using publicly-available spatial transcriptomic data from 10X Genomics (https://www.10xgenomics.com/datasets/xenium-prime-ffpe-neonatal-mouse); the 10X Genomics Cell Segmentation Kit provides cytoskeleton protein information as an individual TIFF file. Segmentation (from the cell boundaries file) and expression (from the feature barcode matrix file) data was acquired from the aforementioned dataset and subject to the standard CytoRipple pipeline. A region of interest was defined using a bounding box based on spatial coordinates corresponding to a section of the developing mouse limb, an area whereby cytoskeleton polarization is known to associate with changes in gene expression profiles. These genes were identified by performing correlation analysis between the Y-component of single-cell polarization vectors and each cell’s associated transcriptome. The top 100 correlated genes, ranked by correlation coefficient, were utilized for pathway enrichment analysis; this analysis was performed using EnrichR^63^ with the gene ontology biological process (GO:BP) library.

Validation of CytoRipple using human colon vimentin polarization was completed using publicly-available spatial proteomic data from *Hickey, Becker, and Nevins et al.*^31^; sample HBM447.MNKW.592 was utilized for these analyses. Cell segmentation and protein intensity TIFF files were acquired from said dataset and subject to the standard CytoRipple pipeline. Comparison to immunohistochemistry staining of vimentin in human colon tissue was completed using publicly-available data from the Human Protein Atlas; sample CAB000080 was utilized for this comparison.

### Processing of Human Melanoma Treated with Immunotherapy

Multiplex ion beam imaging (MIBI) data from our prior work^24^ was subject to the standard CytoRipple pipeline; this prior data contains segmentation, cell type assignment, and protein intensity data for molecular features of interest and cytoskeleton proteins (e.g. vimentin). Patients were stratified by their treatment (combination therapy with nivolumab [anti-PD1] and ipilimumab [anti-CTLA4] or ipilimumab only) and their response to said treatment. Correlation analysis was performed by first subjecting all cytoskeleton dynamic values to percentile transformation (i.e. raw values are transformed into values between 0-100%) and then correlating these percentiles with protein intensity data via Pearson’s method. This correlation analysis was performed in a per-sample manner and in a per-sample and per-cell type manner. For example, a given sample would have correlation values between its spatial proteome data and cytoskeleton dynamics across all cells and for each cell type individually. Correlation coefficients were then aggregated across samples and averaged (via arithmetic mean).

### Regional Analysis of Melanoma with Multi-Cellular Cytoskeleton Dynamics

Spatial regions with similar cytoskeleton dynamics were identified using the Density-Based Spatial Clustering of Applications with Noise (DBSCAN^64^) algorithm; DBSCAN was implemented using scikit-learn^65^. Each cell’s spatial position (X- and Y-spatial coordinates), “curl”, and “divergence” were input into DBSCAN. The latter two inputs were scaled by 1,000x as their default output range is vastly smaller than the numerical range of X- and Y-coordinates; this allows DBSCAN to place greater weight on multi-cellular cytoskeleton dynamics when performing spatial clustering of cells into regions. Only cells that were clustered into regions were utilized for downstream analyses of regions. Regions were then subjected to dimension reduction via principal component analysis (PCA), k-nearest-neighbors (kNN) graph creation, and uniform manifold approximation projection (UMAP^66^); these computations were performed using the Scanpy^67^ package. Unbiased clustering was performed using the Leiden algorithm^68^ with a resolution parameter value of 0.9. Dimension reduction and clustering were performed using information from both multi-cellular cytoskeleton dynamics (“curl” and “divergence”) and protein intensities; the former was given a weight of six compared to the weight of one given to protein intensities to account for the increased number of proteins measured compared to multi-cellular cytoskeleton dynamics and to prioritize information from the cytoskeleton dynamics which was our main feature of interest.

### Trajectory Analysis of Skin and Lung Tumors with Multi-Cellular Cytoskeleton Dynamics

Streams (spatial trajectories) were defined as sequences of cells whose polarization vectors were aligned in a head-to-tail manner (i.e. pointed in the same direction). To identify streams, we subject each cell to the following algorithm: 1) Identify spatially proximal cells (excluding the original cell) within a set radius (this was typically 90µm), 2) Calculate the difference between the original cell’s polarization angle and the polarization angles of spatially proximal cells, 3) Retain proximal cells with polarization angles within π/4 (45 degrees) of the original cell’s polarization angle, 4) If there are no cells stop, otherwise take a step forward with the step’s spatial coordinates and polarization angle set to the average of said values from proximal cells retained from step 3. This algorithm was repeated for each cell up to 25 times and outputs a “stream” of steps in a spatial trajectory with spatial coordinates and multi-omic profiles of associated cells for each step. Null models were built by randomly shuffling spatial coordinates and repeating the aforementioned stream calling process.

Each trajectory step has a “past” (the step before it), “present” (the step itself), and “future” (the step after it). By characterizing trajectory steps via this framework, we can align steps across streams by computing the difference between a step’s present and future from the past and present of all other steps. Intuitively, we are asking two questions: 1) Is this step’s expression profile similar to the “past” of any another step? 2) Is this step’s future expression profile similar to the “present” of any other step? If the answer is yes to both questions, then the identified two steps would appear to be linked, as in they are sequential steps in a common trajectory type. For example, a trajectory step that has high expression of *PD1* with a “future” (subsequent) step with high expression of *TIM3* is likely part of the same “stream” as a trajectory step with a “past” (prior) step with high expression of *PD1* and “present” (current) expression of *TIM3*. We utilized this custom distance algorithm to align trajectory steps across all streams. Dimension reduction (via Potential of Heat-diffusion for Affinity-based Trajectory Embedding [PHATE]^69^) and unbiased clustering (via leiden) can then be performed on this distance matrix. PHATE was chosen instead of UMAP due to its ability to preserve global trajectory structures in high-dimensional data.

The starting cell of each “stream” (spatial trajectories) was defined as the origin, the final step for each “stream” was defined as the terminal. Trajectory steps can then be systematically compared to origins and termini (via their multi-omic expression profiles) to identify their position along an origin-to-termini axis. Alignment of trajectory steps along this axis allows for the characterization of gene and protein expression as a function of their position along spatial trajectories. This analysis was performed across all streams and for each unbiased “stream” cluster. The latter was performed as not all streams in a given tissue will behave the same. For example, some streams may end in immune-rich regions while others may terminate in immunosuppressive regions (e.g. Treg-rich). Correlation analysis of trajectory step expression profiles was performed to cluster of multi-omic parameters (e.g. genes) into modules. The number of modules by computing cluster assignments, via Ward’s algorithm, for a range of cluster numbers and then identifying the number of clusters with the highest silhouette score (i.e. greatest intra-cluster similarity and inter-cluster distance). Module scores were then computed as the average, normalized (via Z-score) expression of genes associated with a given module.

### Quantification and Statistical Analysis

Statistical tests were performed using the Mann-Whitney U test, a non-parametric test that does not assume the data to follow a specific statistical distribution. P-values were corrected for multiple hypothesis testing via the Benjamin-Hochberg method (also known as false discovery rate [FDR]). Correlation was computed using Pearson’s method unless otherwise specified. Clustering was performed using Ward’s method unless otherwise specified.

## Data Availability

All data was publicly available. Intermediate data generated by this manuscript are provided at 10.5281/zenodo.21035128. Video of polarized neutrophils stained for actin was acquired from *Weiner et al.*^23^. Spatial transcriptomic (Xenium) data of the whole mouse pup was acquired from https://www.10xgenomics.com/datasets/xenium-prime-ffpe-neonatal-mouse. Spatial proteomic (CODEX) data of human intestine was acquired from *Hickey, Becker, and Nevins et al.*^31^. Spatial proteomic (MIBI) data of human melanoma treated with immunotherapy was acquired from *Campbell et al.*^24,70,71^. Spatial transcriptomic (Xenium) data of human lung adenocarcinoma was acquired from https://www.10xgenomics.com/datasets/preview-data-ffpe-human-lung-cancer-with-xenium-multimodal-cell-segmentation-1-standard.

## Code Availability

CytoRipple is freely available at https://github.com/danielgchen/CytoRipple as a freely available public, Python, package; tutorial notebooks are available in the same repository. Code for reproduction of the analyses in this manuscript are available at https://github.com/danielgchen/reproducibility_cytoripple.

## Acknowledgements

D.G.C. is supported by the UCLA-Caltech Medical Scientist Training Program and Paul and Daisy Soros Fellowship. J.R.H. acknowledges support from the National Cancer Institute (grant #U54 CA274509-05). A.R. is supported by the Parker Institute for Cancer Immunotherapy (PICI), NIH grants R35 CA197633 and P01 CA168585, the Ressler Family Fund, and the support from Ken and Donna Schultz, Todd and Donna Jones, Robert (Bob) and Mary Jean Rumer, Karen and James Witemyre, and Thomas Stutz through the Jonsson Cancer Center Foundation, and Jonathan Isaacson through the Melanoma Research Foundation. K.M.C. is supported by the V Foundation Gil Nickel Melanoma Research Fellowship, the Parker Institute for Cancer Immunotherapy, and the Melanoma Research Alliance-Ressler Family Fund Young Investigator Award.

## Author Contributions

Conceptualization: D.G.C., M.A., J.R.H., A.R., K.M.C. Data curation: D.G.C., K.M.C. Formal analysis: D.G.C. Funding acquisition: A.R., K.M.C. Investigation: D.G.C., J.S., J.J.P. Methodology: D.G.C. Resources: A.I.L. Software: D.G.C., T.Y., K.M.C. Supervision: K.M.C. Visualization: D.G.C.. Writing - original draft: D.G.C., J.S., J.J.P., K.M.C. Writing - review and editing: D.G.C., T.Y., J.S., J.J.P., A.I.L, M.A., J.R.H., A.R., K.M.C.

## Declaration of Interests

D.G.C. reports consulting fees from Georgiamune. A.R. reports: Honoraria for consulting: Amgen, Bristol-Myers Squibb and Merck; Advisory board and holds stock: Advaxis, Appia, Apricity, Arcus, Compugen, CytomX, Highlight, ImaginAb, ImmPact, ImmuneSensor, Inspirna, Isoplexis, Kite-Gilead, Lutris, MapKure, Merus, PACT, Pluto, RAPT, Synthekine and Tango; Research funding: Agilent and from Bristol-Myers Squibb through Stand Up to Cancer (SU2C); Patent royalties: Arsenal Bio. K.M.C. reports: Shareholder: Geneoscopy LLC, Georgiamune, and AME Therapeutics; Consulting fees: PACT Pharma, Tango Therapeutics, Flagship Labs 81 LLC, the Rare Cancer Research Foundation, Noetik, the Jaime Leandro Foundation, Georgiamune, and AME Therapeutics.

## Supplementary Figures

**Fig. S1:**
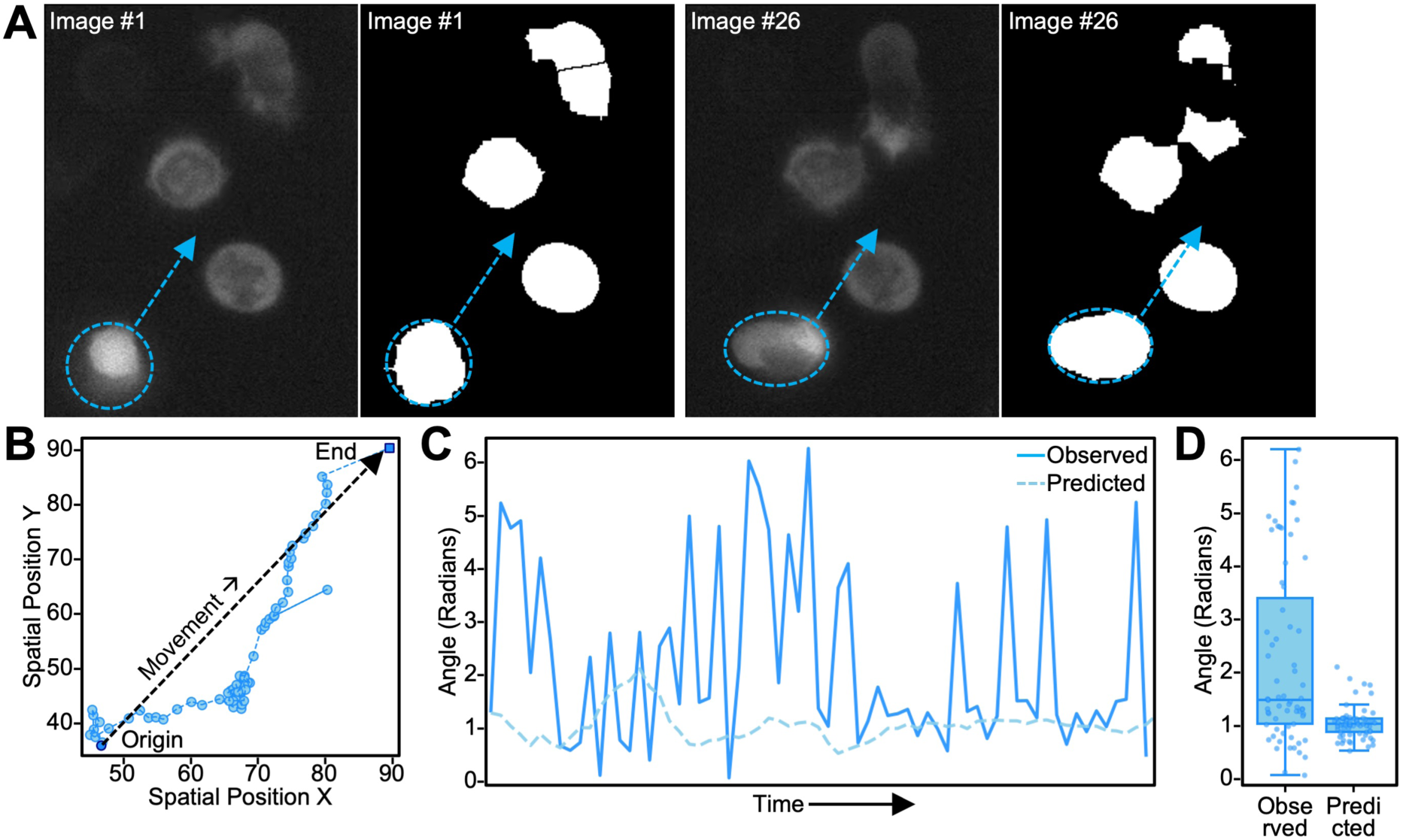
CytoRipple Predictions Align with Observed Trajectory of Neutrophil Polarization. A) Intensity of fluorescently labeled actin (tetramethylrhodamine-actin, left) and the associated ImageJ-based cell segmentation for that given timepoint (right) in chemoattractant stimulated neutrophils. The neutrophil of interest (chosen due to its active movement post chemoattractant stimulation) is circled on the left with a dashed blue line, the arrow indicates the general path it takes throughout time. Two timepoints are shown (image #1 and #26), see labels on the images. B) Scatter plot of the X- and Y-positions of the neutrophil of interest from panel (A). Timepoints are connected by dashed blue lines. The general path the neutrophil takes throughout time is indicated by a thick dashed black arrow. C) Line plot of the observed (solid blue) and CytoRipple predicted (dashed blue) angles of polarization for the neutrophil of interest from panel (A). X-axis represents time, Y-axis represents the angle of polarization in radians. D) Box plot of the observed and predicted (X-axis) angles of polarization (Y-axis). Box plot represents interquartile range (25th to 75th percentiles, IQR) with whiskers representing 1.5 times IQR.

**Fig. S2:**
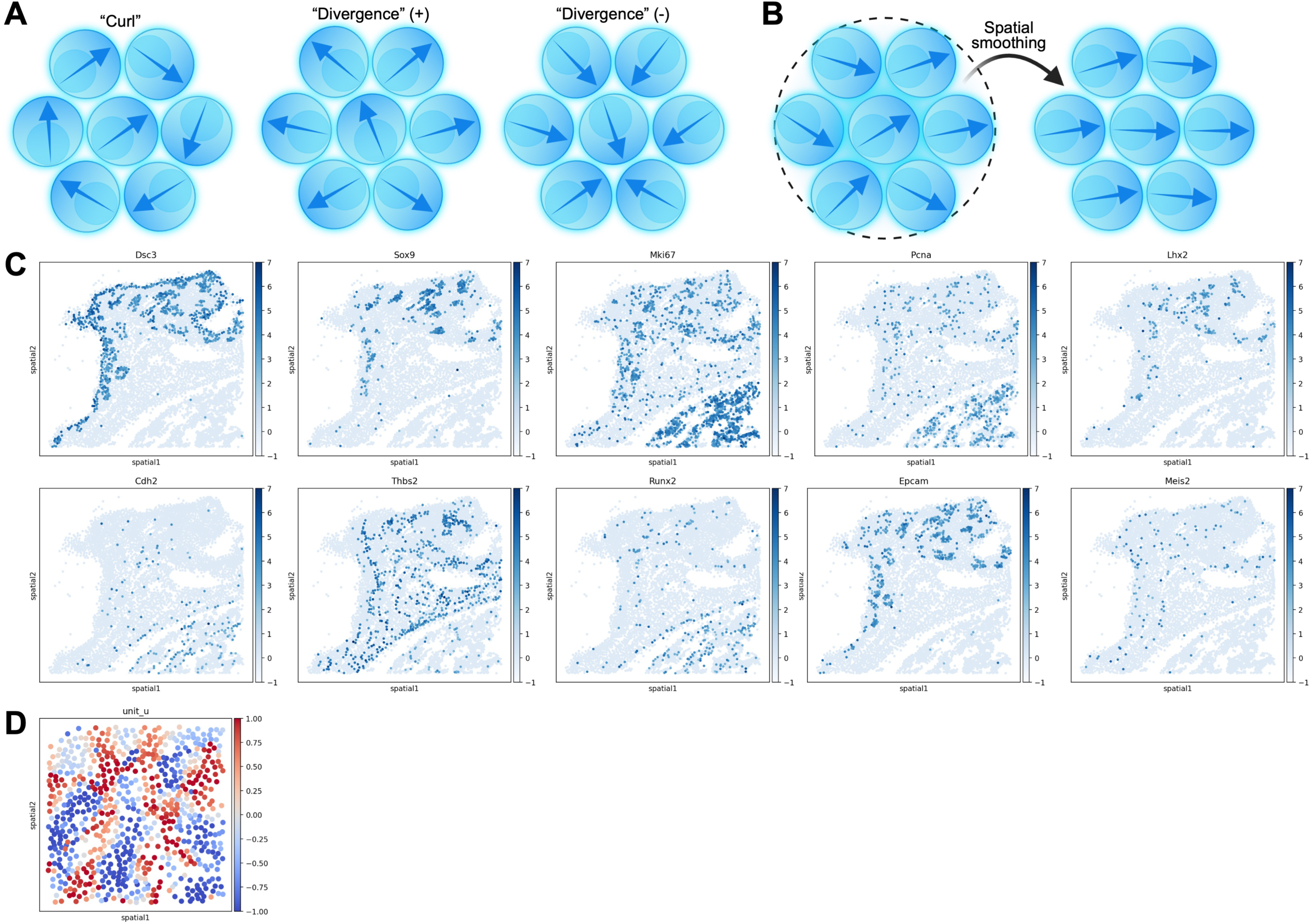
Biological Validation of CytoRipple in the Context of a Developing Mouse Limb. A) Diagram of how different patterns of polarization vectors correspond to different multi-cellular cytoskeleton dynamics (i.e. “curl” and “divergence”). B) Diagram of how spatial smoothing is implemented by CytoRipple. C) Spatial plots of the expression of limb development marker genes in the developing mouse limb from Fig. 2D (see legends on the right). D) Spatial plots of the X-component of polarization angles in the developing mouse limb’s region of interest from Fig. 2D (see legend on the right).

**Fig. S3:**
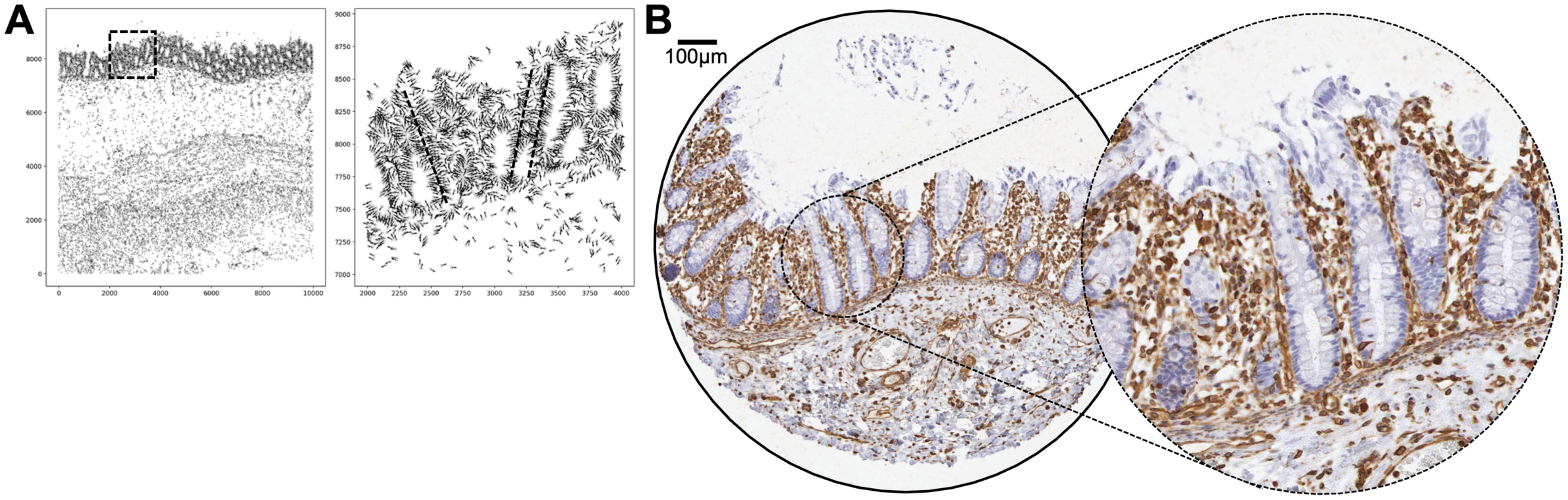
Biological Validation of CytoRipple in the Context of Human Colon Crypts. A) Spatial plot of the human colon (large intestine) and a region of interest (chosen for its capture of multiple linear crypts). The region of interest is denoted in the original image by a dashed blue outline. Each dot represents a cell, and arrows represent polarization vectors. B) Representative staining of vimentin in human colon by immunohistochemistry. Image was acquired from the Human Protein Atlas.

**Fig. S4:**
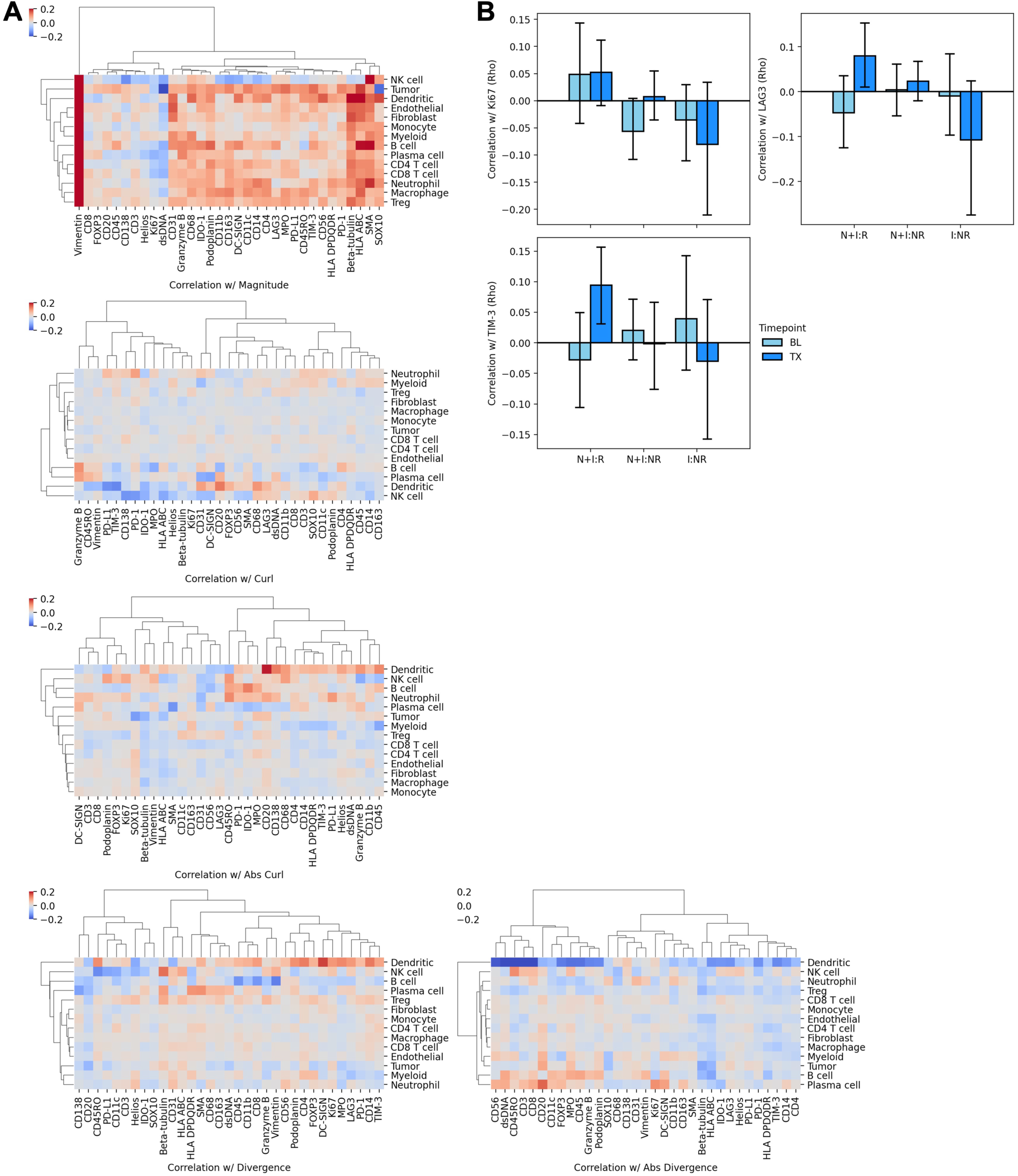
Cell Type Specific Protein Expression Associations with Cytoskeleton Dynamics in Patients with Melanoma Treated with Immunotherapy. A) Heatmaps of the correlation (Pearson’s method) between cytoskeleton dynamics and protein expression (columns) for each cell type (rows). Color represents average correlation coefficient (rho) across samples (see legend on the right). B) Bar plots of the coefficients of correlation (rho) between divergence (a cytoskeleton dynamic) and CD8^+^ T cell protein expression (Y-axis) as stratified by patient treatment and response (X-axis). Color represents timepoint: pre-treatment in dark blue and post-treatment in light blue (see legend on the left). P-values were calculated using the Mann-Whitney U test. Bar heights represent arithmetic mean and error bars represent 95% confidence interval.

**Fig. S5:**
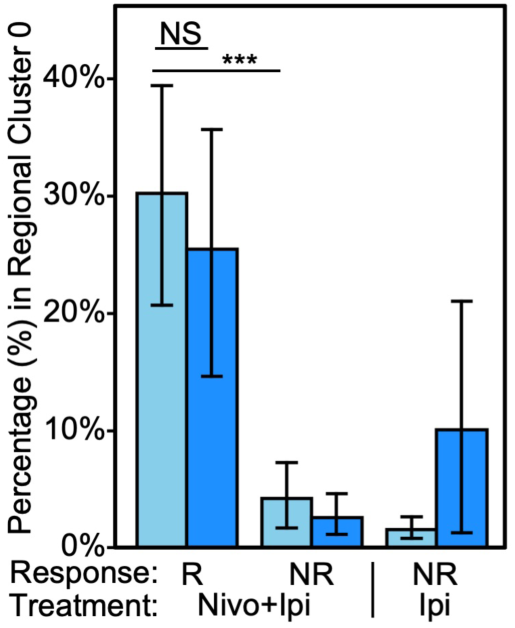
Regional Cluster 0 is Overrepresented in Responders to Immunotherapy. Bar plot of the percentage (%) of cells in each sample that are in regional cluster 0 (Y-axis) as stratified by patient treatment and response (X-axis). Color represents timepoint: pre-treatment in dark blue and post-treatment in light blue (see legend on the right). P-values were calculated using the Mann-Whitney U test. Bar heights represent arithmetic mean and error bars represent 95% confidence interval.

**Fig. S6:**
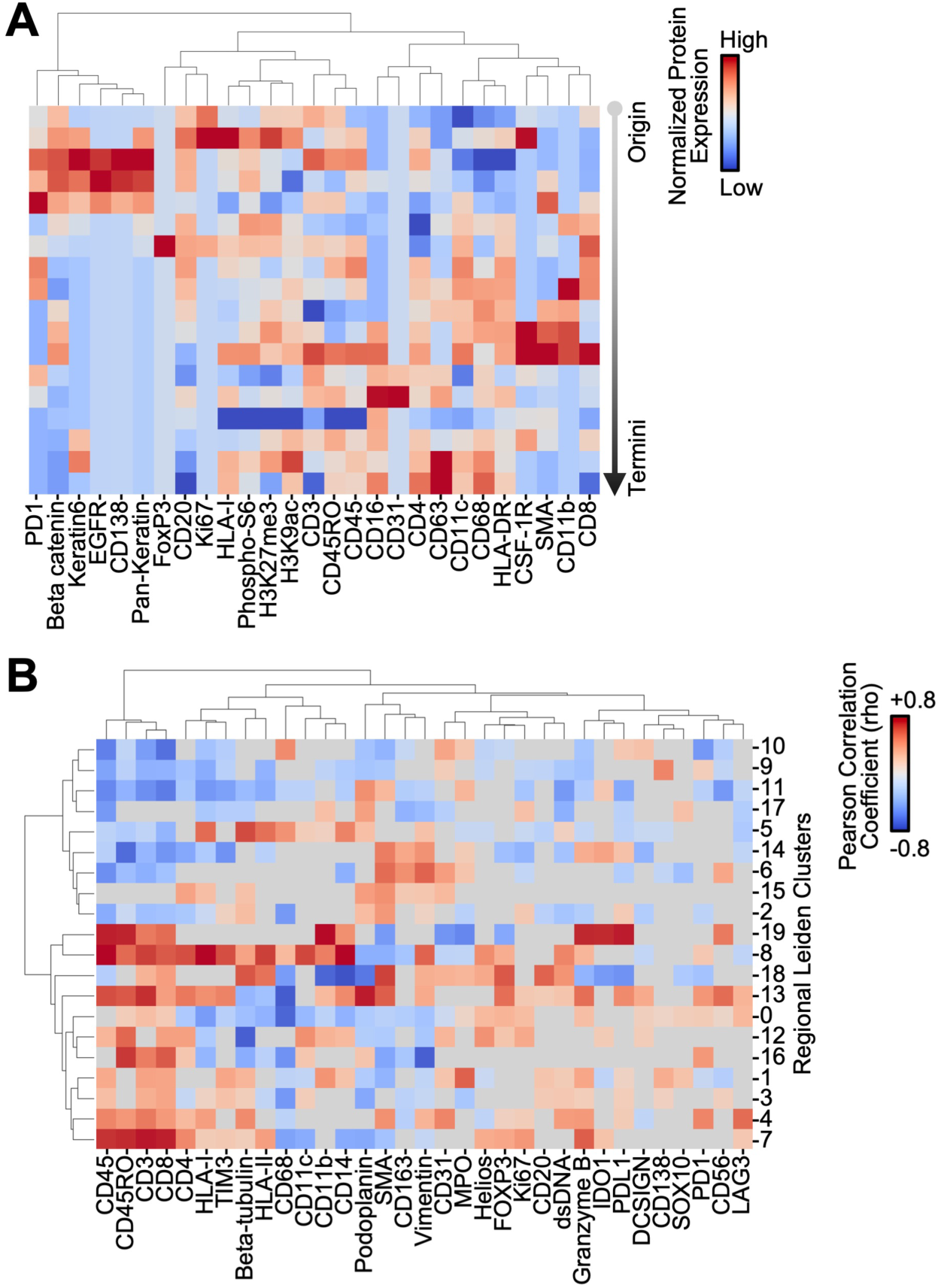
Heterogeneous Spatial Trajectories Exist in a Biopsy of Human Melanoma. A) Heatmap of the expression of proteins (columns) across trajectory steps (rows) in the representative stream from Fig. 5A. Color represents expression magnitude (see legend on the right). B) Heatmap of the expression of proteins (columns) in each trajectory cluster (rows). Color represents expression magnitude (see legend on the right).

**Fig. S7:**
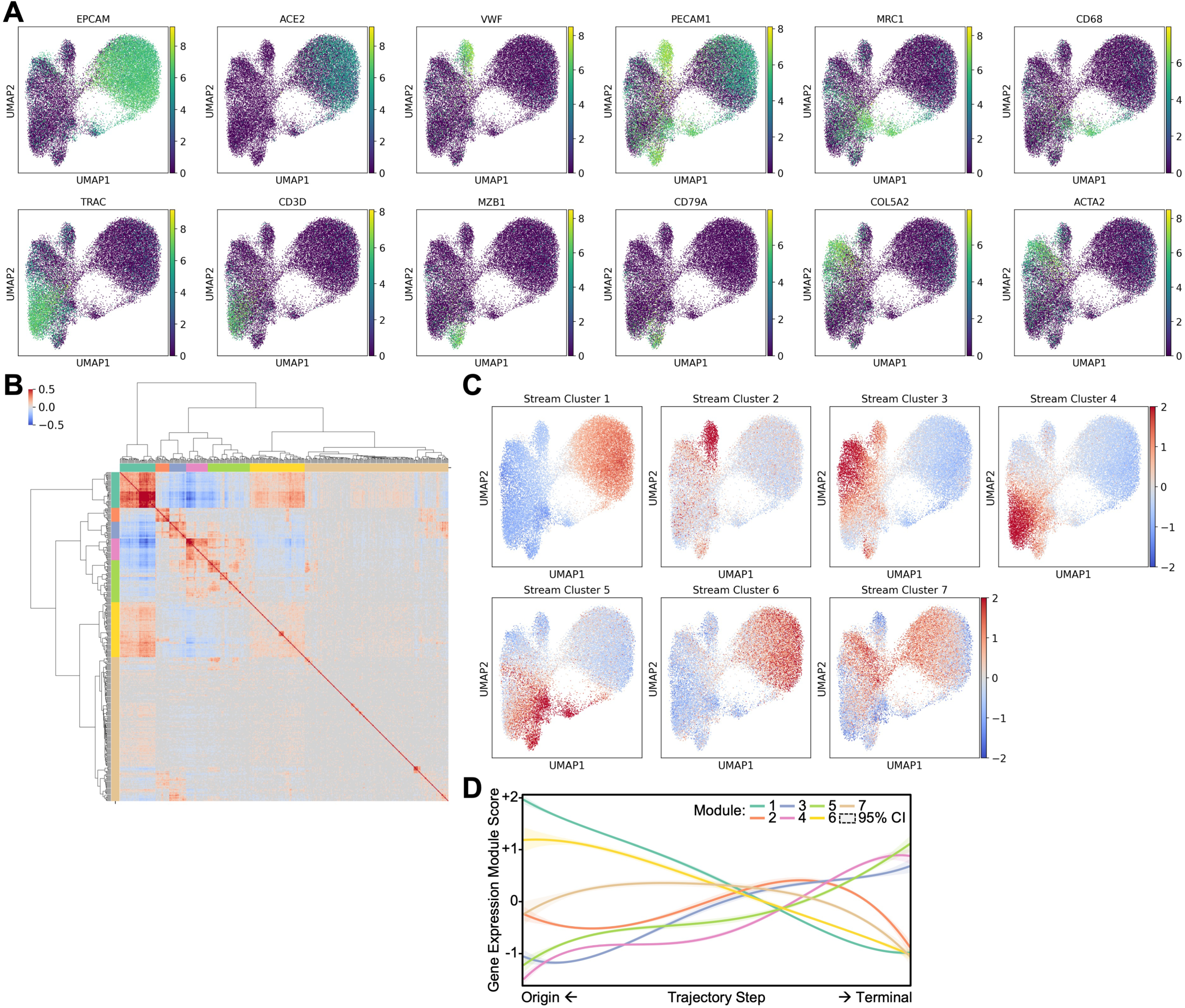
A Central Spatial Trajectory Exists in a Biopsy of Human Lung Adenocarcinoma. A) UMAP of single cell transcriptomes from a biopsy of human lung adenocarcinoma. Dots are colored by their expression of cell type marker genes (e.g. *VWF* for endothelial cells, see legends on the right). B) Heatmap of the correlation matrix of trajectory step expression matrices. Color represents correlation coefficient (computed via Pearson’s method, see legend on the upper left). Significant (p<0.05) correlations are presented; non-significant correlations are represented by a light gray color. Correlation coefficients of each gene are clustered via Ward’s algorithm. C) UMAP of single cell transcriptomes from a biopsy of human lung adenocarcinoma. Color represents normalized expression (Z-score) of gene modules (see legends on the right). Module identities are labeled above each plot. D) Line plot of normalized expression (Z-score, Y-axis) of gene modules presented in panels (B) and (C) across trajectory steps (X-axis). Trajectory steps are ordered by their gene expression distance from cells that are origins of streams (similar cells on the left) and distance from cells that are termini of streams (similar cells on the right). Color represents module identity and shaded areas represent 95% confidence interval (see legend on the upper right).

## Supplementary Tables

Table S1: Correlation (Pearson’s method) between gene expression and the Y-vector of cells in the developing limb of a whole mouse pup.

Table S2: Pathway enrichment analysis (from GO:BP) of genes positively correlated with the Y-vector of cells in the developing limb of a whole mouse pup.

Table S3: Differentially expressed genes for cells associated with streams versus all other cells in a biopsy of human lung adenocarcinoma.

Table S4: Correlation (Pearson’s method) between gene expression profiles of streams in a biopsy of human lung adenocarcinoma; correlation coefficient (rho).

Table S5: Correlation (Pearson’s method) between gene expression profiles of streams in a biopsy of human lung adenocarcinoma; correlation significance (p-value).

